# A new mutant mouse model lacking mitochondrial-associated CB_1_ receptor

**DOI:** 10.1101/2020.03.30.009472

**Authors:** Antonio C. Pagano Zottola, Edgar Soria-Gomez, Itziar Bonilla-del-Río, Carolina Muguruza, Geoffrey Terral, Laurie M. Robin, José F. Oliveira da Cruz, Bastien Redon, Thierry Lesté-Lasserre, Tarson Tolentino-Cortes, Nagore Puente, Gabriel Barreda-Gómez, Francis Chaouloff, Luis F. Callado, Pedro Grandes, Giovanni Marsicano, Luigi Bellocchio

**Author notes:** Correspondence to: Luigi Bellocchio, Giovanni Marsicano. These authors share first authorship (Antonio C. Pagano Zottola, Edgar Soria-Gomez). These authors share senior authorship (Giovanni Marsicano, Luigi Bellocchio).

## Abstract

The idea that the effects of drugs largely depend on subcellular target location is emerging as a novel predictive factor of their beneficial or adverse outcomes. G protein-coupled type-1 cannabinoid receptors (CB_1_) are regulators of several brain functions as well as the main targets of cannabinoid-based medicines.

Besides their classical location at plasma membranes, CB_1_ receptors are present at different locations within cells, including in association to mitochondrial membranes (mtCB_1_). Here we report the generation and characterization of a mutant mouse line, which lack mtCB_1_ receptors.

## INTRODUCTION

As a classical seven transmembrane G protein coupled receptor, CB_1_ is associated with plasma membranes (pmCB_1_), where it mediates communication between extracellular and intracellular signaling (Ibsen et al., 2017) and it regulates neurotransmitter and neuropeptide release (Busquets-Garcia et al., 2018). However, recent evidence from our and others’ laboratories indicates that functional CB_1_ receptors can be also found intracellular, particularly in association with mitochondrial membranes (mtCB_1_) (Gutierrez-Rodríguez et al., 2018; Hebert-Chatelain et al., 2014; Koch et al., 2015; Mendizabal-Zubiaga et al., 2016). We recently showed that hippocampal mtCB_1_ receptors are responsible for the amnesic effects of cannabinoids *via* alteration of bioenergetic processes (Hebert-Chatelain et al., 2016). Indeed, it is well known that impaired mitochondrial functions negatively affect synaptic plasticity and lead to cognitive and locomotor behavioral abnormalities (Kann and Kovacs, 2007; Rangaraju et al., 2014). Here we generate and characterize a new mutant mouse model to better study the functions of mitochondrial associated CB_1_ receptors.

## METHODS

### Mice

Experiments were approved by the Committee on Animal Health and Care of INSERM and the French Ministry of Agriculture and Forestry (authorization number 3306369). Mice were maintained under standard conditions (food and water *ad libitum*; 12h/12h light/dark cycle, light on 7 a.m.; experiments were performed between 9 a.m. and 5 p.m.). DN22-*CB*_*1*_-KI mice were generated using a flox-stop strategy as previously described (Ruehle et al., 2013). Stop cassette was excised *via* a CRE deleter mouse strain and knock-in mice were maintained over a C57BL6/N background for several generations before experiments.

### Quantitative real-time PCR (q-PCR)

Samples from WT, DN22*-CB*_*1*_-KI and *CB*_*1*_-KO mice were homogenized in Tri-reagent (Euromedex, France) and RNA was isolated using a standard chloroform/isopropanol protocol (Chomczynski and Sacchi, 1987). RNA was processed and analyzed following an adaptation of published methods (Bustin et al., 2009). cDNA was synthesized from 1 μg of total RNA using Maxima Reverse Transcriptase (Thermo Scientific) and primed with oligo-dT primers (Thermo Scientific) and random primers (Thermo Scientific). qPCR was perfomed using a LightCycler® 480 Real-Time PCR System (Roche). qPCR reactions were done in duplicate for each sample, using transcript-specific primers, cDNA (4 ng) and LightCycler 480 SYBR Green I Master (Roche) in a final volume of 10 μl. The PCR data were exported and analyzed in an informatics tool (Gene Expression Analysis Software Environment) developed at the NeuroCentre Magendie. For the determination of the reference gene, the Genorm method was used (Livak and Schmittgen, 2001). Relative expression analysis was corrected for PCR efficiency and normalized against two reference genes. Valosin containing protein (Vcp) and succinate dehydrogenase complex subunit (Sdha) genes were used as reference genes for Amygdala. Glyceraldehyde-3-phosphate dehydrogenase (Gapdh) and succinate dehydrogenase complex subunit (Sdha) genes were used as reference genes for Hippocampus. Succinate dehydrogenase complex subunit (Sdha) and tyrosine 3 mono oxygenase/tryptophan 5 mono oxygenase activation protein zeta (Ywhaz) genes were used as reference genes for Anterior Olfactory Nucleus. Succinate dehydrogenase complex subunit (Sdha) and tubulin alpha 4 a (Tuba4a) genes were used as reference genes for Prefrontal cortex. Actin beta (Actb) and tubulin alpha 4 a (Tuba4a) genes were used as reference genes for Caudate putamen. Glyceraldehyde-3-phosphate dehydrogenase (Gapdh) and peptidylprolyl isomerase A (Ppia) genes were used as reference genes for Hypothalamus. The relative level of expression was calculated using the comparative (2^-ΔCT^) method (Livak and Schmittgen, 2001). Primers sequences are reported in the following table:

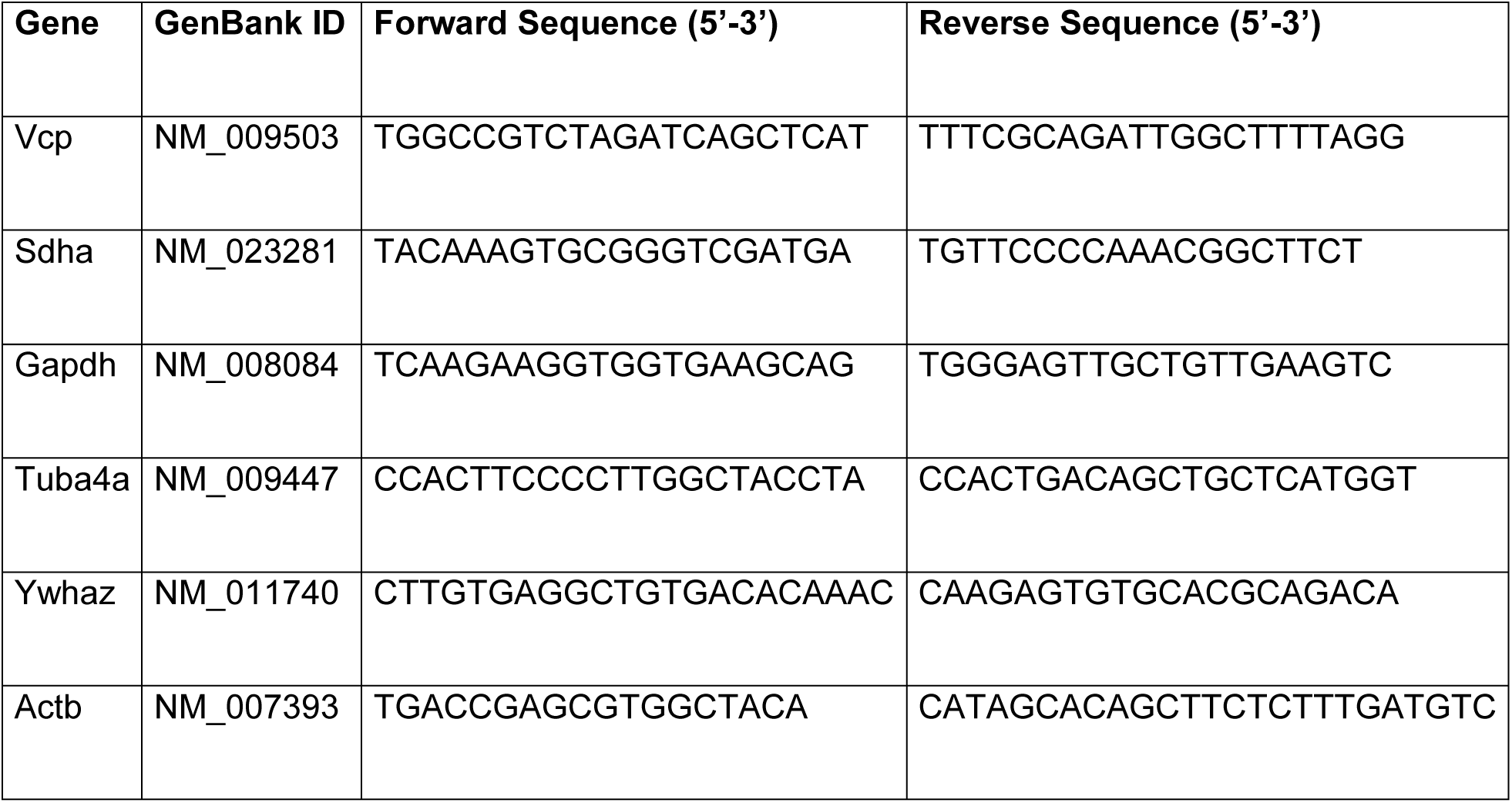
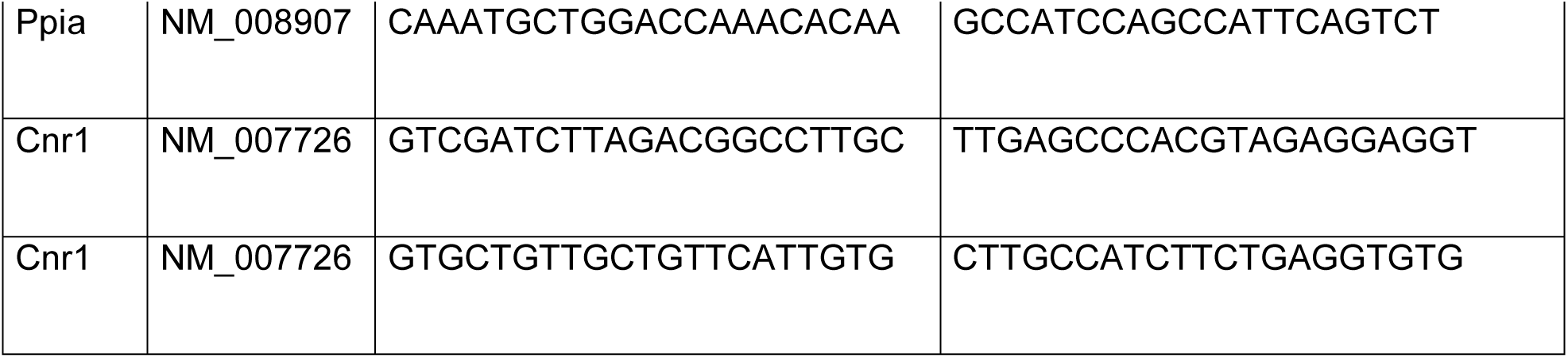

### Mitochondrial respiration

Mitochondrial respiration in substantia nigra (SN) extracts was measured as previously described (Hebert-Chatelain et al., 2016) with some modification. Briefly, SN was rapidly dissected from mouse brain using the coronal brain matrix(Vallee et al., 2014) and was homogenized in 450 *μ*l of Miro5 buffer (Makrecka-Kuka, et al., Biomolecules 2015) using a Politron homogenizer (11.000 rpm 3-5 sec). After brief centrifugation the supernatant was treated with saponin at a final concentration of 12.5 *μ* g/ml. Respiration analyses were carried out using a 2K Oroboros device (Makrecka-Kuka et al., 2015). 100 µl of lysate were put in each chamber and complex I-dependent respiration was triggered by adding malate (0.5 mM), pyruvate (5 mM) and glutamate (10 mM) (MPG) (Makrecka-Kuka et al., 2015). Then we applied DMSO or WIN55.212-2 (Sigma Aldrich, FRANCE) at final concentration of 1µM and 5 minutes after we injected 1.25 mM ADP. Each measure of Oxygen consumption rate (OCR) in ADP condition was normalized to the MPG values before ADP injection and the effect of WIN was expressed as change respect to vehicle conditions. The assay was performed in duplicate for each sample. Only samples with a ratio of ADP/MPG superior to 1.5 were retained for the analyses.

In a previous set of experiments we validated the quality of mitochondria in the preparation by measuring the ADP/MPG ratio and then we added the complex II substrate Succinate at the final concentration of 10mM, 10µM Cytochrome C, to chek mitochondrial membrane integrity, and finally Rotenone (0.5µM) and Antimycin A (2.5µM) inhibitors of complex I and III, respectively (Makrecka-Kuka et al., 2015) (**Table 1**).

**Table 1.**
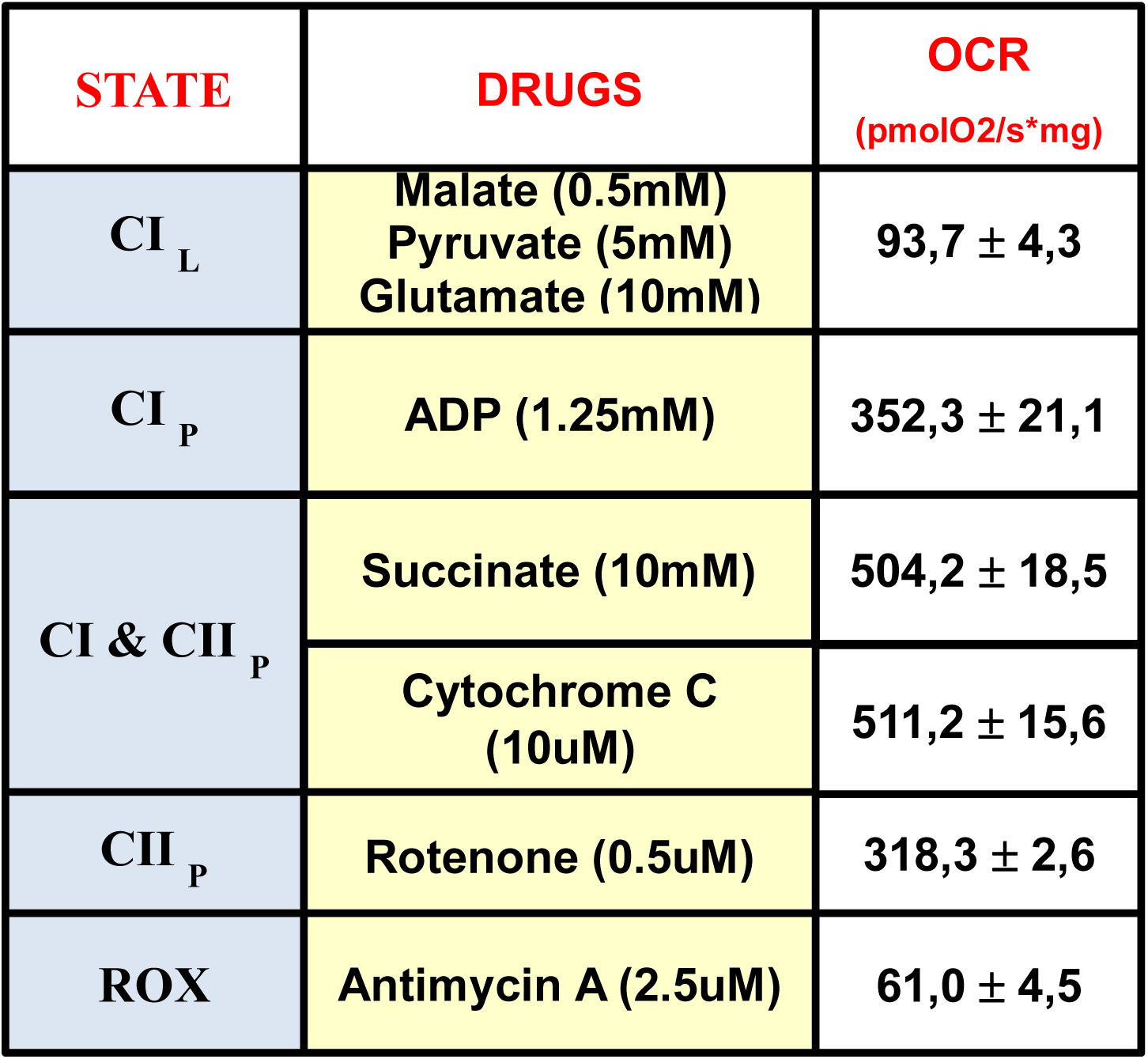
Characterization of Mitochondrial respiration in SNr extracts. CI _L_, ComplexI Leak; CI _P_, ComplexI OXPHOS; CI & CII _P_, ComplexI and ComplexII OXPHOS; CII _P_, ComplexII OXPHOS; ROX, Residual Oxygen Consumption.

| STATE | DRUGS | OCR<br>(pmolO<sub>2</sub>/s*mg) |
| --- | --- | --- |
| CI<sub>L</sub> | Malate (0.5mM)<br>Pyruvate (5mM)<br>Glutamate (10mM) | 93,7 ± 4,3 |
| CI<sub>P</sub> | ADP (1.25mM) | 352,3 ± 21,1 |
| CI & CII<sub>P</sub> | Succinate (10mM) | 504,2 ± 18,5 |
|  | Cytochrome C<br>(10uM) | 511,2 ± 15,6 |
| CII<sub>P</sub> | Rotenone (0.5uM) | 318,3 ± 2,6 |
| ROX | Antimycin A (2.5uM) | 61,0 ± 4,5 |

### [^35^S]GTPγS binding assay on brain homogenates

Brains from 8 mice of each genotype (DN22-*CB*_*1*_-KI, WT and *CB*_*1*_-KO) were obtained. Midbrain, cortex and hippocampus were dissected and immediately stored at − 70 °C until assay. For each brain region, tissue samples from each genotype were pooled to obtain the enriched fractions of plasma membranes. Tissue samples were homogenized using an a Teflon-glass grinder (IKA Labortechnik, Satufen, Germany) at 1.500 rpm (10 up-and-down strokes) in 30 volumes of homogenization buffer (1 mM EGTA, 3 mM MgCl2, 1 mM DTT, and 50 mM Tris-HCl, pH 7.4) supplemented with 0.25 M sucrose. The crude homogenate was centrifuged for 5 minutes at 1,000 x g at 4 °C and the supernatant layer was re-centrifuged for 10 minutes at 40,000 x g (4°C). The resultant pellet (P2 fraction) was washed twice in 10 and 5 volumes of homogenization buffer respectively, and re-centrifuged in similar conditions. Protein content was measured according to Bradford’s method using Bovine Albumine Serum (BSA) as standard. Samples were aliquoted in order to have a protein content of 1 mg and then centrifuged in a benchtop centrifuge (EBA 12 R, Hettich Instruments, Tuttlingen, Germany) at highest speed (14,000 rpm) during 15 minutes at 4 °C. The supernatant layer was carefully discarded and the pellets stored at -70 °C until assay. The tissue samples of the nine experimental conditions (3 genotypes and 3 brain regions, 1 pool for each region-genotype) were processed in parallel on the same day. The day of the experiment the membrane pellets were defrosted (4 °C), thawed and re-suspended in 11 ml of incubation buffer containing 1 mM EGTA, 3 mM MgCl2, 100 mM NaCl, and 50 mM Tris-HCl, pH 7.4, reaching a final protein concentration of 0.09 mg/ml approximately. The real final protein content was measured after the experiment according to Bradford’s method.

WIN55.213-2 stimulated [^35^S]GTPγS binding assays were carried out in a final volume of 250 μl in 96 well plates, containing 1 mM EGTA, 3 mM MgCl2, 100 mM NaCl, 0.2 mM DTT, 50 μM GDP, 50 mM Tris–HCl at pH 7.4 and 0.5 nM [^35^S]GTPγS. Stimulation curves were carried out by incubating increasing concentrations WIN (10-^12^-10-^4^ M); 9 concentrations by duplicate; three independent experiments. The incubation was started by addition of the membrane suspension (18 μg of membrane proteins per well) and was performed at 30 °C for 120 minutes with shaking (450 rpm). Incubations were terminated by rapid filtration under vacuum (1450 FilterMate Harvester, PerkinElmer) through GF/C glass fiber filters (Printed Filtermat A) pre-soaked in ice-cold incubation buffer. The filters were then rinsed three times with 300 μl ice-cold incubation buffer, air dried (20°C, 120 minutes), and counted for radioactivity (4 minutes) by liquid scintillation spectrometry using a MicroBeta TriLux counter (PerkinElmer). Non-specific binding of the radioligand was defined as the remaining [^35^S]GTPγS binding in the presence of 10 μM unlabelled GTPγS, and the basal binding, as the signal in the absence of agonist. The pharmacological parameters of the stimulation curves of the [^35^S]GTPγS binding, the maximal effect (Emax) and the concentration of the drug that determines the half maximal effect (EC50), were obtained by non-linear analysis using GraphPad Prism™ software version 5.0. The points fit to a concentration-response curve (standard slope). The pharmacological parameters Emax and EC50 are expressed as means ± SEM. The statistical comparison of the data sets was performed in GraphPad Prism™ software version 5.0, by a co-analysis of the curves.

### [^35^S]GTPγS and [^3^H]CP55,940 binding assay on brain slices

[^35^S]GTPγS (1250 Ci/mmol) and [^3^H]CP55,940 (149 Ci/mmol) were purchased from PerkinElmer (Boston MA, USA). The [^14^C] and [^3^H] standards were supplied by American Radiolabelled Chemicals (St. Louis, MO, USA). DL-dithiothreitol (DTT), guanosine-5’-diphosphate (GDP), and guanosine-5’-γ-3-thiotriphosphate (GTPγS) were acquired from Sigma-Aldrich (St. Louis, MO, USA). WIN 55,212-2 was purchased from Tocris Bioscience (Bristol, UK). All other chemicals were obtained from standard sources and were of the highest purity commercially available.

In order to test the ability of CB1 receptor to stimulate GTPγS conversion after WIN administration or to bind the CP55,940 agonist, 5-6 WT and DN22-*CB1*-KI mice, and 1 *CB1*-KO mouse as control, were sacrificed by cervical dislocation and their brain rapidly frozen at -80°C. 20µm slices were cut with a cryostat and mounted on superfrost (Thermo Scientific, FRANCE) slides.

#### [^35^S]GTPγS binding assay

Brain sections were thawed for 15 min and then incubated in 50 mM Tris-HCl buffer with 3 mM MgCl_2_, 0.2 mM EGTA, 100 mM NaCl, 2 mM GDP and 1 mM DTT (pH = 7.4) for 20 min at room temperature. Afterwards, the slices were incubated with 0.04 nM [^35^S]GTPγS in absence and in presence of WIN 55,212-2 μM (1 and 10 μM) for 2 h at 30°C. The non-specific binding was determined with 10 μM GTPγS. Finally, the sections were washed twice in 50 mM Tris-HCl (pH =7.4) for 15 min at 4°C, dried and exposed to a Kodak Biomax MR film with ^14^C standards. The films were scanned and quantified by transforming the optical densities into nCi/mg using the ^14^C standards (NIH-IMAGE, Bethesda, MA, USA). The background and the non-specific densities were subtracted. The percentages of stimulation were calculated from the basal and agonist-stimulated [^35^S]GTPγS binding densities according to the formula (stimulated x 100/basal) - 100.

#### [^3^H]CP55,940 binding assay

Tissue sections were dried and incubated in 50 mM Tris-HCl buffer containing 1% of BSA (pH 7.4) for 30 min at room temperature. Later, the brain sections were incubated again in the same buffer supplemented with 3 nM [^3^H]CP55,940 for 2h at 37°C. Non-specific binding was determined with 10 μM WIN 55,212-2. Finally, sections were washed in ice-cold 50 mM Tris-HCl buffer supplemented with 1% BSA followed by dipping in distilled water at 4°C. After drying, the brain slides were exposed to a radiation sensitive film for 21 days at 4°C together with tritium standards. The films were scanned and quantified by transforming the optical densities into nCi/mg using the tritium standards (NIH-IMAGE, Bethesda, MA, USA). The background and the non-specific densities were subtracted to determine the specific binding.

### Behavioral tests

#### Wire hang test

Wild type and DN22-*CB*_*1*_-KI mice were tested for their muscular strength in the wire hang test. All tests were conducted during the dark phase of the nycthemeral cycle under dim red light. Each cage, housing one mouse, was placed on a bench in a room adjacent to the one housing the mice. Thereafter, each mouse was removed from its cage, placed on the cage grid, the latter being then slowly inverted as to be suspended 90 cm above a big cage filled with polystyrene beads. The latency to fall was then recorded individually.

#### Basal locomotion

The basal locomotor activity of WT and DN22-*CB*_*1*_-KI mice was measure during seven days using the TSE PhenoMaster system (TSE Systems GmbH). Mice were placed individually in a plexiglass cage [45cm (length) X 34cm (width) X 20cm (height)] surrounded with the ActiMot module containing IR light beams recognizing locomotor activity. Food and water were available *ad libitum*.

### Immuno-electron microscopy

*CB*_*1*_-KO, DN22-*CB*_*1*_-KI, and respective WT littermates mice (n = 3 per genotype) were deeply anesthetized by intraperitoneal injection of ketamine/xylazine (80/10 mg/kg body weight) and were transcardially perfused at room temperature (RT, 20-25°C) with phosphate buffered saline (0.1 M PBS, pH 7.4) for 20 s, followed by the fixative solution made up of 4% formaldehyde (freshly depolymerized from paraformaldehyde), 0.2% picric acid, and 0.1% glutaraldehyde in phosphate buffer (0.1 M PB, pH 7.4) for 10-15 min. Then, brains were removed from the skull and post-fixed in the fixative solution for approximately 1 week at 4°C. Afterwards, brains were stored at 4°C in 1:10 diluted fixative solution until used.

### Pre-embedding silver-intensified immunogold method

Coronal vibrosections at the level of mesoencephalum containing the substantia nigra *pars reticulata* were cut at 50 □m and collected in 0.1 M phosphate buffer (PB) (pH 7.4) with 0.1% sodium azide at RT. Sections were preincubated in a blocking solution of 10% BSA, 0.1% sodium azide, and 0.02% saponin prepared in Tris-HCl buffered saline (TBS 1X, pH 7.4) for 30 min at RT. Then, sections were incubated with a primary goat anti-CB_1_ antibody binding to a 31 amino acids sequence of the C-terminus (CB_1_ C-ter^31^; 2 µg/ml; Cat. N.: CB_1_-Go-Af450-1; Frontier Institute; Japan) in the blocking solution but with 0.004% saponin on a shaker for 1 day at RT. After several washes in 1% BSA/TBS, tissue sections were incubated in a secondary anti-goat 1.4 nm gold-labeled Immunoglobulin-G antibody (Fab’ fragment, 1:100, Nanoprobes Inc.) in 1% BSA/TBS with 0.004% saponin on a shaker for 4 h at RT. Thereafter, the tissue was washed in 1% BSA/TBS overnight at 4°C and post-fixed in 1% glutaraldehyde in TBS for 10 min at RT. Following washes in double distilled water, gold particles were silver-intensified with a HQ Silver kit (Nanoprobes Inc., Yaphank, NY, USA) for about 12 min in the dark and then washed in 0.1 M PB (pH 7.4). Stained sections were osmicated (1% osmium tetroxide, OsO_4_, in 0.1 M PB pH 7.4, 20 min), dehydrated in graded alcohols to propylene oxide, and plastic-embedded flat in Epon 812. Ultrathin sections of 50 nm were collected on mesh nickel grids, stained with 2.5% lead citrate for 20 min, and examined in a Philips EM208S electron microscope. Tissue preparations were photographed by using a digital camera coupled to the electron microscope. Figure compositions were made at 600 dots per inch (dpi). Labeling and minor adjustments in contrast and brightness were made using Adobe Photoshop (CS, Adobe Systems, San Jose, CA, USA).

### Semi-quantification of mtCB_1_ receptor immunostaining using immunogold method

2-3 of 50 µm-thick sections containing the substantia nigra *pars reticulata* and hippocampus from each animal genotype (n=3 each) showing good and reproducible silver-intensified gold particles were cut at 50 nm. Electron micrographs were taken from grids with similar labeling intensity indicating that selected areas were at the same depth. To avoid false negatives, only ultrathin sections in the first 1.5 µm from the surface were examined. To determine the proportion of CB_1_-positive mitochondria, the percentages of CB_1_ immunopositive mitochondria in each mice were calculated taking into account only particles (at least one) on mitochondrial membrane segments far away from other membranes (distance ≥ 80 nm). Image-J (version 1.36) was used to measure this distance. On the other hand, the proportion of CB_1_ in mitochondria versus total CB_1_ signal was analyzed in each image. Graphs and statistical analyses were performed using GraphPad software (version 5.0).

### Electrophysiology

#### Hippocampal depolarization-induced suppression of inhibition

Male DN22-*CB*_*1*_-KI mice and their WT littermates were sacrificed by dislocation and the brain was immediately immerged in ice-cold oxygenated cutting solution containing in mM: 180 Sucrose, 26 NaHCO3, 12 MgSO4, 11 Glucose, 2.5 KCL, 1.25 NaH2PO4, and 0.2 CaCl2, oxygenated with 95% O2-5% CO2 ≈300mOsm. Parasagittal hippocampal slices (300μm thick) were obtained using a vibratome (VT1200S, Leica, Germany) and transferred for 30min into a 34°C bath of oxygenated ACSF containing in mM: 123 NaCl, 26 NaHCO3, 11 Glucose, 2.5 KCL, 2.5 CaCl2, 1.3 MgCl2, 1.25 NaH2PO4 ≈305 mOsm. After a minimum of 1h recovery at room temperature (22-25°C), slices were transferred to a recording chamber in ACSF at 32°C. Whole-cell recordings of IPSCs were made using a MultiClamp 700B amplifier (Molecular devices, UK) in CA1 pyramidal neurons voltage clamped at −70mV with a pipette (3-5 MΩ) containing in mM: 130 KCl, 10 HEPES, 1 EGTA, 2 MGCl2, 0.3 CaCl2, 7 Phosphocreatin, 3 Mg-ATP, 0.3 Na-GTP; pH = 7.2; 290mOsm. Evoked IPSCs were performed by a monopolar stimulating patch pipette filled with ACSF in stratum radiatum in presence of NMDA and AMPA/Kainate receptor antagonists (50μM D-APV and 10μM NBQX). DSIs were performed by depolarizing pyramidal neurons from −70mV to 0mV for 3 s. DSIs’ magnitude were measured as the average of 3 DSIs with 2min apart and represented the percentage of change between the mean of the 5 consecutive IPSCs preceding the depolarization and the first three IPSCs following the depolarization, with IPSCs evoked every 3 s. Currents were filtered at 4kHz by a Digidata 1440A (Molecular devices, UK) and were analyzed using either Clampfit software (pClamp10).

### Statistical analyses

All graphs and statistical analyses were performed using GraphPad software (version 5.0 or 6.0). Results were expressed as means of independent data points ± s.e.m. Data were analyzed using unpaired Student’s *t*-test or ANOVA (followed by Bonferroni’s *post hoc* test), as appropriate. Post hoc significances were expressed as follow: * p<0.05, ** p<0.01, *** p<0.001.

## RESULTS AND DISCUSSION

The potential implication of different subcellular pools of CB_1_ receptors on the regulation of brain processes, by both endogenous cannabinoids and exogenously administered ligands, is still unknown.

In order to solve this issue, we developed a new mutant knock-in mouse line. We previously showed that a mutant version of the CB_1_ protein lacking the first 22 amino acids (called DN22-CB_1_) is neither anatomically or functionally associated to mitochondrial membranes, nevertheless maintaining its other functions (Hebert-Chatelain et al., 2016). Here, we generated DN22-*CB*_*1*_-KI mice, in which the wild-type *CB*_*1*_ receptor gene is replaced by the coding sequence of DN22-CB_1_ protein (**Figure 1A**). When compared to their wild-type DN22-*CB*_*1*_-WT littermates, DN22-*CB*_*1*_-KI mice presented the same amount of CB_1_ mRNA expression in different brain regions, measured by qPCR (**Figure 1B**), although a slight non-significant decrease of CB_1_ protein was observed by immunofluorescence in certain brain regions (**Figure 1C**). Counting of gold particles in immuno-electron microscopy experiments indicated that the total amount of CB_1_ receptor protein in the substantia nigra pars reticulate (SNr) is not significantly different between DN22-*CB*_*1*_-KI and DN22-*CB*_*1*_-WT littermates (**Figures 1D and 1E**). Similarly, the levels of CB_1_ receptor associated to plasma membrane in the same brain region were not impacted by the mutation (**Figures 1D and 1F**). Strikingly, however, the amount of mitochondrial-associated CB_1_ receptors (mtCB1) and of CB_1_-positive mitochondria over total number of these organelles were significantly lower in DN22-*CB*_*1*_-KI mice as compared to WT littermates, reaching levels undistinguishable from global *CB*_*1*_-KO mice (**Figures 1D, 1G and 1H**).

**Figure 1.**
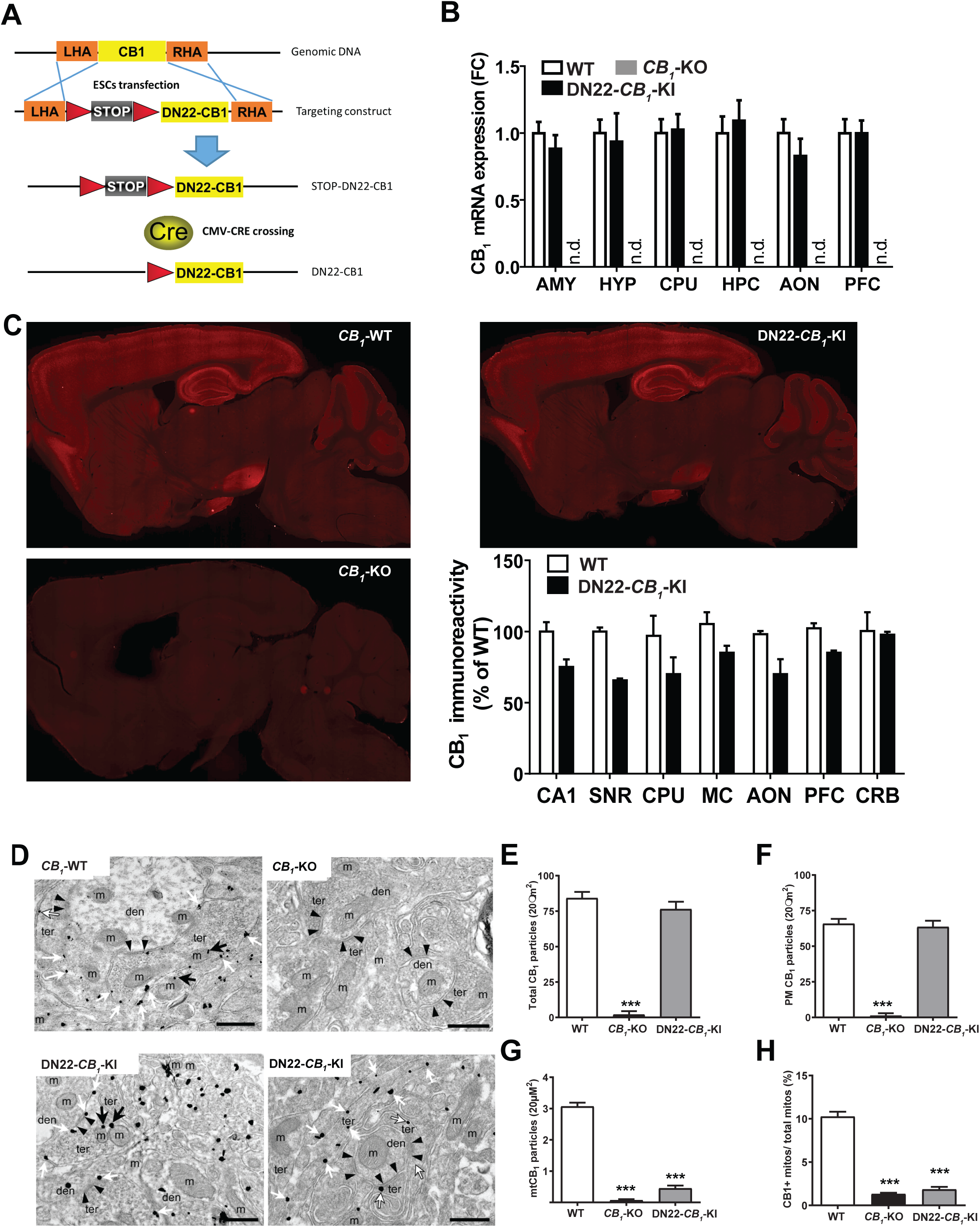
**(A)** Schematic representation of the strategy employed for the generation of DN22-CB_1_-KI mouse line. LHA: left homology arm; RHA, right homology arm. **(B)** qPCR analysis of CB_1_ mRNA in different brain regions from WT, DN22-CB1-KI and CB1-KO mice. AMY, amygdala; HYP, hypothalamus; CPU, caudate putamen; HPC, hippocampus; AON, anterior olfactory nucleus; PFC, prefrontal cortex. **(C)** Immunofluorescence detection of CB_1_ receptor in the brain of WT vs DN22-CB_1_-KI mice (CB_1_-KO mice are shown as antibody negative control). CA1, hippocampal region; SNR, substantia nigra reticulata; CPU, caudate putamen; MC, motor cortex; AON, anterior olfactory nucleus; PFC, prefrontal cortex; CRB, cerebellum. **(D)** Immunogold detection of CB_1_ receptor coupled to electron microscopy in WT, *CB*_*1*_-KO and DN22-*CB*_*1*_-KI mice in SNr terminals and relative quantification of total gold particles and mitochondrial (mtCB_1_) particles. White arrows, plasma membrane gold particles; black arrows, mitochondrial gold particles; scale bar = 500 nm. Den, dendrites; ter, synaptic terminal, m, mitochondrion. **(E and F)** Relative quantification of total gold particles and plasma membrane (PM-CB_1_) particles. **(G and H)** Relative quantification of mitochondrial gold (mtCB_1_) particles and gold-positive mitochondria over total.

To test the functionality of CB_1_ receptor-dependent G protein activation, we performed [^35^S]GTPγ binding assays in cortical, hippocampal and midbrain extracts from WT, *CB*_*1*_-KO, and DN22-*CB*_*1*_-KI mice in the presence of the cannabinoid agonist WIN55,212 (WIN). The results clearly indicated that the DN22-CB_1_ mutant protein is as efficient in triggering G protein activation as its wild-type cognate (**Figure 2A**). These results were also confirmed in other brain regions by *in situ* [^35^S]GTPγ binding assays in brain sections (**Figures 2B and 2C**). Moreover, auto radiographic analysis of [^3H^]CP55.940 cannabinoid agonist binding in the same setup, indicated no differences between the two genotypes in different brain regions (**Figure 2D and 2E**). Thus, DN22-CB_1_ mutation does not seem to impact on the global ability of CB_1_ receptors to bind cannabinoid ligands and subsequently activate G-proteins release.

**Figure 2.**
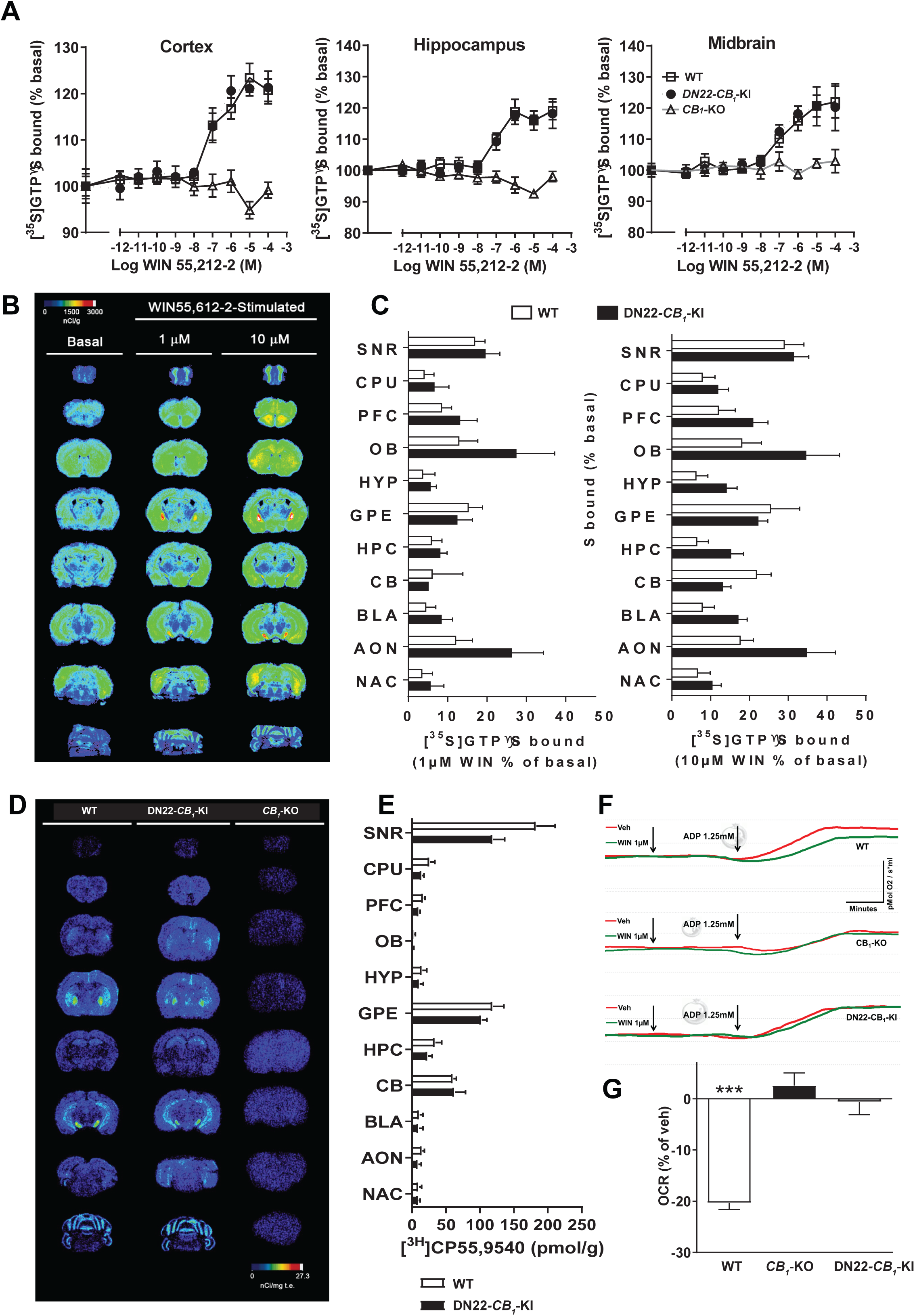
**(A)** Dose–response curves of the effect of WIN 55,212-2 on [^35^S]GTPγS binding in membranes isolated from cortex, hippocampus and midbrain of WT, *CB*_*1*_-KO and DN22-*CB*_*1*_-KI mice. **(B)** Representative autoradiograms of brain sections from basal, 1 µM and 10 µM WIN55,212-2-stimulated [35S]GTPγS binding in wt mice. **(C)** Relative quantification of WIN-mediated [35S]GTPγS over basal level at 1µM (left) and 10µM (right) doses in wt and DN22-CB_1_-KI mice. SNR, substantia nigra reticulata; CPU, caudate putamen; PFC, prefrontal cortex; OB, olfactory bulb; HYP, hypothalamus; GPE, external globus pallidus; HPC, hippocampus; CB, cerebellum; BLA, basolateral amygdala; AON, anterior olfactory nucleus; NAC, nucleus accumbens. **(D)** Representative autoradiograms of brain sections from WT, DN22-CB_1_-KI and CB_1_-KO mice incubated with [^3H^] CP55.940. **(E)** Relative quantification of bound [^3H^] CP55.940 in WT, DN22-CB_1_-KI and CB_1_-KO slices. SNR, substantia nigra reticulata; CPU, caudate putamen; PFC, prefrontal cortex; OB, olfactory bulb; HYP, hypothalamus; GPE, external globus pallidus; HPC, hippocampus; CB, cerebellum; BLA, basolateral amygdala; AON, anterior olfactory nucleus; NAC, nucleus accumbens. **(F)** Representative trace of mitochondrial respiration in a substantia nigra preparation from WT, DN22-CB_1_-KI and CB_1_-KO mice. Substrates are malate, pyruvate and glutamate. The inhibitory effect of WIN was observed when the respiration was coupled to ATP synthesis via the addition of Adenosine diphosphate (ADP, see **Table 1**). **(G)** The effect of WIN (1µM) on mitochondrial respiration in SNr homogenates from WT, *CB*_*1*_-KO, and DN22-*CB*_*1*_-KI mice. Data are expressed as percentage of vehicle values.

Oxygen consumption assays in SN homogenates revealed that WIN lowers ADP-stimulated mitochondrial respiration in WT but not in *CB*_*1*_-KO mice (**Figures 2F** and **2G**), indicating a specific CB_1_-mediated control of mitochondrial activity in the SN. Notably, this effect was abolished in SN homogenates from DN22-*CB*_*1*_-KI mice, showing its dependency on mtCB_1_ receptors (**Figure 2F and 2G**).

DN22-*CB*_*1*_-KI mutant strain is viable, fertile and presented normal body weight, muscular strength and locomotor activity (**Figures 3A and 3B**). Interestingly, DN22-*CB*_*1*_-KI mice did not show any alteration in voluntary running wheel activity (data not shown), where global *CB*_*1*_-KO mice display a clear impairment (Dubreucq et al., 2013). Conversely, amnesic effect of cannabinoids in an L-maze object recognition task (Puighermanal et al., 2009), which is mediated by mtCB_1_ receptor signaling in the hippocampus (Hebert-Chatelain et al., 2016), was fully abolished in DN22-*CB*_*1*_-KI mice (data not shown). Previous evidence suggests that mtCB_1_ receptors might be partially responsible for electrophysiological depolarization-induced suppression of inhibition in the hippocampus (DSI, Benard et al. 2012), which is well-known to depend on endocannabinoid signaling mobilization (Zou and Kumar, 2018). Interestingly, DN22-*CB*_*1*_-KI mice displayed a DSI of significantly smaller amplitude as compared to WT littermates (**Figures 3C**), confirming the participation of mtCB_1_ in this form of synaptic plasticity (Benard et al. 2012).

**Figure 3.**
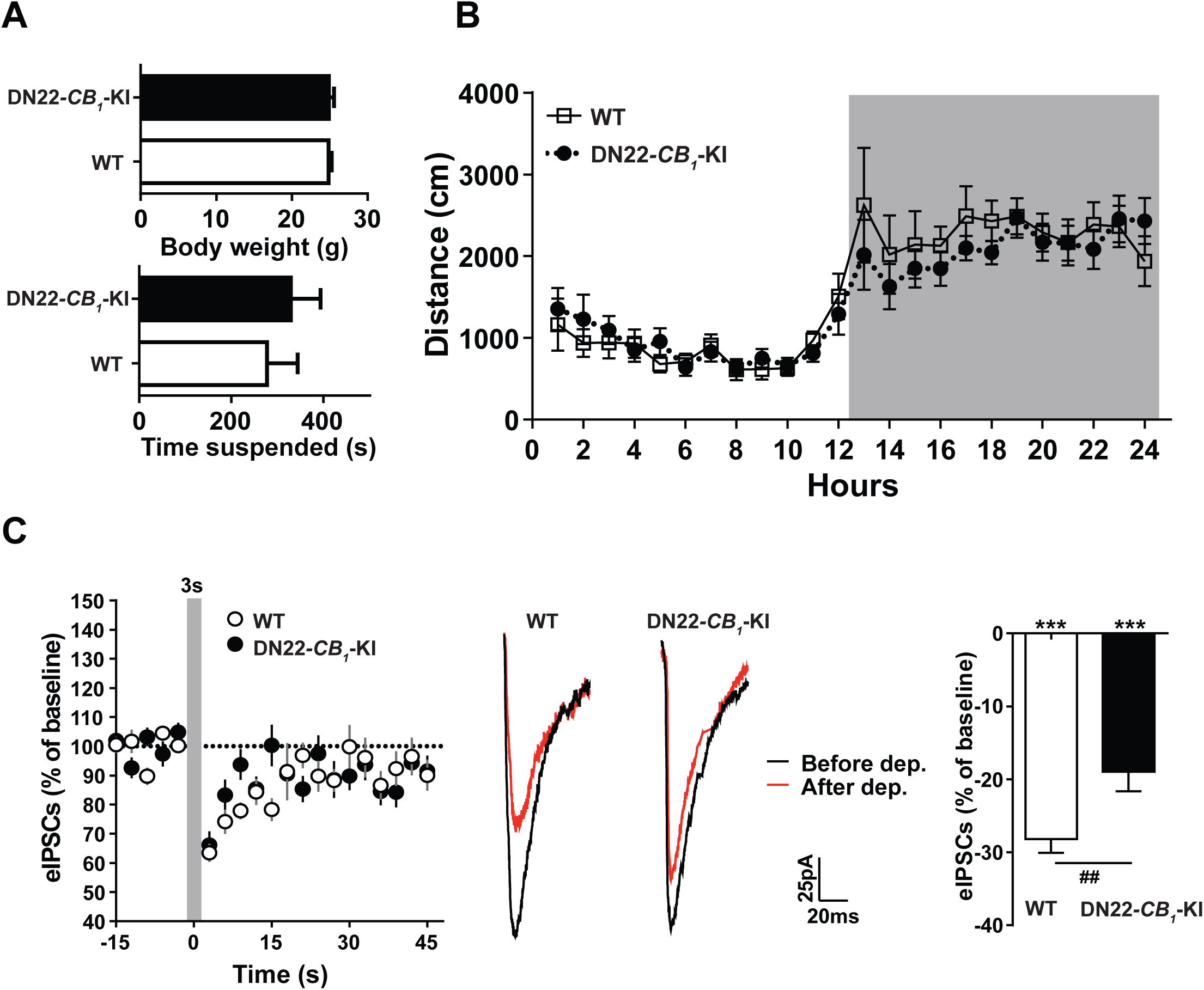
**(A)** Body weight (top panel) and muscular strength (measured by grid suspension test, bottom panel) of WT and DN22-CB_1_-KI mice. **(B)** Daily home cage locomotor activity of WT and DN22-CB_1_-KI mice. Grey box indicate the night phase. **(C)** Left, time course plot showing eIPSCs amplitude before and after 3s depolarization (−70 to 0 mV grey line) in hippocampal slices obtained from WT and DN22-CB_1_-KI mice. Right, representative traces of eIPSCs before and after 3s depolarization (−70 to 0 mV) in WT and DN22-CB_1_-KI mice and averaged reduction of eIPSCs amplitude respect to baseline recorded during 3 sweeps after depolarization.

Altogether, these observations indicate that the constitutive deletion of the first 22 aminoacids of the CB_1_ protein in mice specifically impact the effects of cannabinoids involving mitochondrial activity, but leave other functions of CB_1_ receptors unchanged or reduced when mtCB_1_ receptors are involved. Thus, the DN22-*CB*_*1*_-KI mouse strain represents the *bona fide* tool to study the role of mitochondrial-associated CB1 receptor in the regulation of ECS-mediated brain functions.

## ACKNOWLEDGMENTS

We thank Delphine Gonzales, Nathalie Aubailly, and all the personnel of the Animal Facility of the NeuroCentre Magendie for mouse care. We also thank all the members of the Marsicano lab for useful discussions, Virginie Morales for invaluable help with administrative work. We thank the Histology and Biochemistry platforms of the NeuroCentre Magendie, as well as the Bordeaux Imaging Center (BIC) for help in the experiments. We thank Etienne Hebert-Chatelain and Su Melser for helpful suggestions regarding the mitochondrial respiration and Zhe Zhao for helping in image acquisition. We also thank Daniela Cota and Manuel Guzman for their useful and critical reading on the manuscript. This work was supported by INSERM (to G.M. and L.B.), EU–FP7 (PAINCAGE, HEALTH-603191 to G.M.), European Research Council (Endofood, ERC–2010–StG–260515; CannaPreg, ERC-2014-PoC-640923, Micabra to G.M.), Fondation pour la Recherche Medicale (DRM20101220445 to G.M. and ARF20140129235 to L.B.). Human Frontiers Science Program (to G.M.), Region Aquitaine (to G.M.), French State/Agence Nationale de la Recherche (LABEX BRAIN ANR-10-LABX-43 to G.M and JCJC mitoCB1-FAT to LB.); Fyssen Foundation, Ikerbasque (The Basque Foundation for Science), MINECO (Ministerio de Economía y Competitividad) PGC2018-093990-A-I00, to E.S-G.

